# Tonic Inhibition is Abolished in GABA_A_ Receptor γ2R43Q Knock-in Mice with Absence Epilepsy and Febrile Seizures

**DOI:** 10.1101/155556

**Authors:** Kile P. Mangan, Wyatt B. Potter, Aaron B. Nelson, Steve Petrou, Stephen M. Johnson, Avtar Roopra, Chiara Cirelli, Mathew V. Jones

## Abstract

The γ2R43Q GABA_A_ receptor mutation confers absence epilepsy in humans, and γ2R43Q knock-in mice (RQ) display absence seizures and generalized spike-and-wave discharges reminiscent of their human counterparts. Previous work on several rodent models led to the conclusion that elevated tonic inhibition in thalamic neurons is necessary and sufficient to produce typical absence epilepsy. In contrast, here we used patch-clamp electrophysiology in brain slices to show that RQ mice entirely lack tonic inhibition in principal cells of layer II/III somatosensory cortex and ventrobasal thalamus. Additionally, protein quantification and multielectrode electrophysiology show that the mutation interferes with trafficking of GABA_A_ receptor subunits involved in generating tonic currents, leading to increased cortical firing and decreased thalamic bursting rates. Together with previous work, our results suggest that an optimum level of tonic inhibition is required for normal thalamocortical function, such that deviations in either direction away from this optimum enhance susceptibility to absence seizures.

---

Several human epilepsies have been traced to mutations in the GABA_A_ receptor^1,2^, a pentameric transmembrane protein containing an integral chloride ion channel that regulates action potential generation via shunting or hyperpolarization. The mutation that has received the most study is an arginine-to-glutamine substitution at position 43 of the γ2 subunit (γ2R43Q)^3^. Human patients harboring the γ2R43Q mutation present symptoms from a variety of epileptic phenotypes, the most common being Childhood Absence Epilepsy (CAE) and febrile seizures^3^. γ2R43Q knock-in mice (RQ) display absence seizures and generalized EEG spike-and-wave discharges (SWDs) reminiscent of their human counterparts^4^ (see Sup. Fig.). Absence seizures consist of brief losses of consciousness typically lasting 2-15 s, along with bilateral, synchronous 3-Hz spike-and-wave discharges (SWDs)^5^. Human patients with the γ2R43Q mutation show evidence of a hyperexcitable cortex compared to unaffected family members, displaying increased intracortical excitability, decreased intracortical inhibition and increased facilitation in response to paired-pulse stimulation^6^. These findings support the hypothesis that a hyperexcitable cortical condition is thought to contribute to SWDs in these patients^7^. A similar “cortical focus theory” for absence seizures was proposed after SWD generation was localized to the somatosensory cortex in a different mouse model^8,9^. The exact origin of SWDs in γ2R43Q human patients has not been identified.

The functional effects of the γ2R43Q mutation have been studied in heterologous expression systems (oocytes, HEK293 and COS7 cells), but have led to conflicting results. On one hand, the γ2R43Q mutation has been shown to alter receptor function by slowing receptor deactivation, enhancing desensitization, and reducing benzodiazapine sensitivity^10,11^. However, others have observed little effect on receptor function^12^. In contrast, several studies agree that the mutation alters GABA_A_ receptor assembly, trafficking or surface expression^12,13,14,15,16,17^. Interestingly, this mutation in the γ2 subunit appears to also affect trafficking of other subunits including α1, α3, β2, β3, and α5^13,16,17^. The α5 subunit participates in extrasynaptic tonic inhibition in cortex, and is decreased by the γ2R43Q mutation^16^. Whether the γ2R43Q mutation also affects trafficking of the δ subunit, which contributes to thalamic tonic inhibition, is not yet known.

Tonic inhibition has recently been linked to SWD generation and absence seizures^18^. Multiple rodent models of absence epilepsy (GAERS, stargazer, lethargic, tottering) display increases in thalamic inhibitory tonic current, and selective activation of this current produces SWDs and absence seizures in rats^18,19^. To understand how altered thalamic inhibitory tonic currents could produce SWDs, we must consider the anatomy and functional connectivity of neurons in the thalamocortical network.

Thalamic relay neurons can fire in distinct ‘tonic’ and ‘burst’ modes. The tonic firing mode occurs when the membrane is steadily depolarized, and consists of classical sodium channel-dependent action potentials. The burst firing mode, in comparison, occurs when the membrane is hyperpolarized such that T-type voltage-gated calcium channels are allowed to deinactivate. A subsequent depolarization then results in a high frequency burst of sodium channel-dependent action potentials riding atop a calcium channel-dependent plateau potential. Thus, increased hyperpolarizing tonic inhibition may shift thalamic relay neurons into the burst firing mode^20^, which may increase the drive onto GABAergic thalamic reticular nucleus (TRN) neurons. In turn, TRN neurons transmit hyperpolarizing IPSPs back onto thalamic relay neurons, further promoting relay neuron burst firing. This reverberation between relay and TRN neurons is critical for sustaining SWDs^21,22,23^. Indeed, even in studies supporting a cortical origin of SWDs^9^, seizure activity spread to the thalamus within a few hundred milliseconds, consistent with the idea that robust absence seizures are a product of the full thalamocortical network^22,24^.

Here we show, using thalamocortical slices, that tonic inhibition is abolished in layer II/III neurons of somatosensory cortex and relay neurons of ventrobasal thalamus of RQ mice. Through Western blotting and voltage-clamp electrophysiology, we show that the loss of tonic inhibition is accompanied by altered expression or trafficking of the GABA_A_ receptor subunits responsible for mediating tonic currents in these areas. Using multielectrode arrays, we further show that loss of tonic inhibition increases cortical firing rates, but decreases bursting throughout the thalamocortical circuit, consistent with a depolarization of thalamic relay neurons that shifts them away from the burst firing mode. Selective pharmacological blockade of cortical tonic current in wild type (RR) slices also increases cortical firing rates, paralleling the increased cortical firing in RQ slices, and consistent with the increased cortical excitability observed in γ2R43Q human patients. Together these results suggest that the combined loss of cortical and thalamic tonic inhibition in RQ mice enhances susceptibility to absence seizures.

## RESULTS

### Synaptic inhibition is reduced in RQ thalamus and cortex

The RQ mutation hinders GABA_A_ receptor assembly, trafficking and surface expression^12,13,14,15,16,17^, and decreases cortical mIPSC amplitude in RQ mice^4^. Our analysis of mIPSCs corroborated the latter finding, showing a decrease (31%) in mIPSC amplitude in somatosensory cortical layer II/III neurons (pA: mean ± SEM, N; RR: 26.2 ± 2.9, 9; RQ: 18.1 ± 1.4, 8, p<0.05; Fig. 1a & 1b) and also a decrease (34%) in thalamic relay neurons (RR: 41.1 ± 4.8, 9; RQ: 27.1 ± 2.1, 7, p<0.05; Fig. 1d & 1e). mIPSC frequency was unaffected in both areas (Fig. 1c & 1f). Weighted decay time-constants for cortical layer II/III neurons are not different for RQ (ms: mean ± SEM, N; 24.0 ± 1.6, 4) compared to RR (19.0 ± 2.2, 5), but are increased for RQ thalamic neurons compared to RR (RR: 8.5 ± 0.8, 7; RQ: 13.8 ± 0.6, 5, p<0.001) (data not shown).

**Figure 1.**
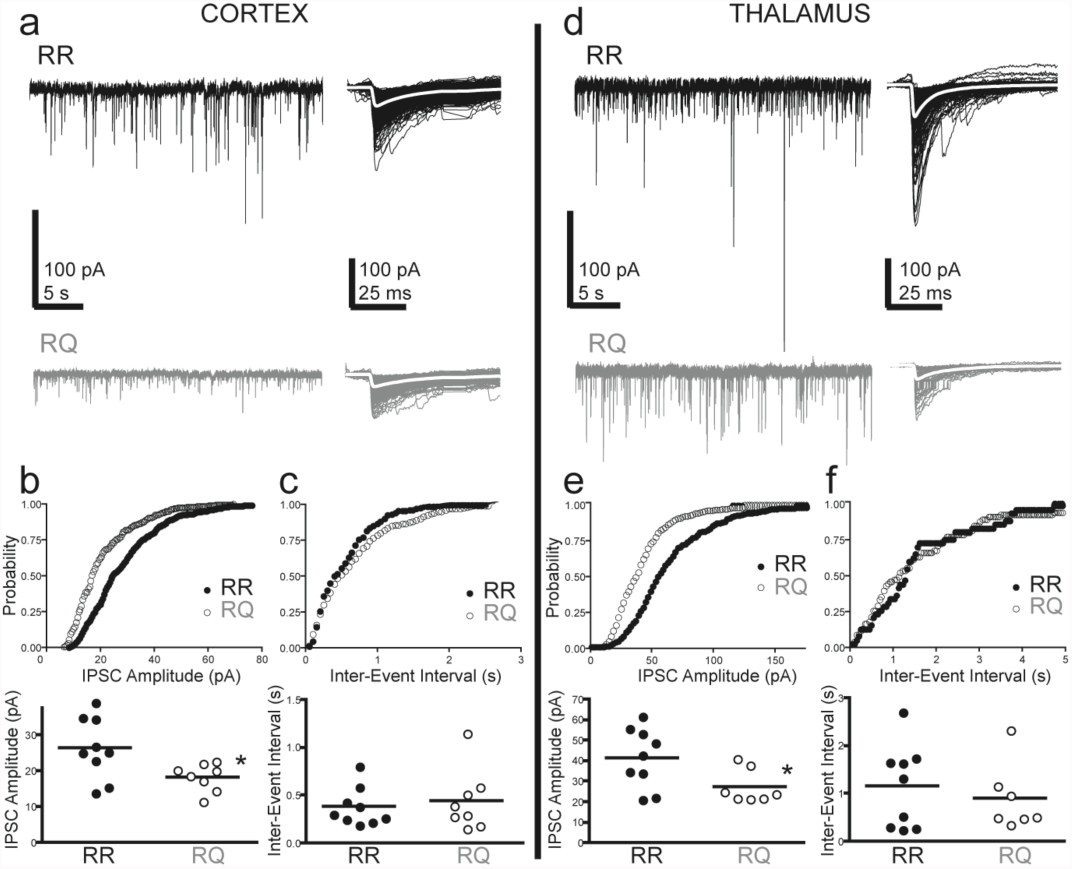
mIPSC amplitude is decreased in RQ slices. A) Example voltage-clamp trace (left) and corresponding miniature inhibitory postsynaptic currents (mIPSCs) (right) for a wild type (RR) (top: black) and a mutant (RQ) (bottom: grey) cortical layer II/III pyramidal cell. Overlayed white traces are the average mIPSCs. B) Cummulative amplitude distributions (top) and median mIPSC amplitudes (bottom) for cortical cells, showing a reduction in mIPSC amplitude for RQ compared to RR (p<0.05, asterisk). Bars represent the mean of medians. C) Cummulative interevent interval (IEI) distribution (top) and medians (bottom) for cortical neurons, showing no difference in RQ compared to RR. D-F) Same as A-C, but for thalamic relay neurons in RR and RQ slices.

### GABAergic tonic inhibition is abolished in RQ cortical and thalamic neurons

Although reductions in synaptic inhibition are a potential mechanism for hyperexceitability and absence epilepsy in the RQ mice, the γ2R43Q mutation may affect other processes as well. For example, based on studies in transfected cultured neurons, Eugène et al.^16^ proposed that this mutation may contribute to absence epilepsy by reducing tonic inhibition. Therefore, to directly test this hypothesis in an animal model, we examined tonic inhibition in slices from RR and RQ knock-in mice. Using whole cell voltage clamp recordings we found that, whereas RR neurons exhibit a substantial inhibitory tonic current, this current was entirely abolished in RQ mutant somatosensory cortical layer II/III neurons (pA: mean ± SEM, N; RR: 5.8 ± 0.6, 5; RQ: -1.2 ± 1.6, 4, p<0.05; Fig. 2a & 2b), as well as in thalamic relay neurons (RR 10.6 ± 3.6, 9; RQ: 0.6 ± 0.7, 6, p<0.05; Fig. 2c & 2d).

**Figure 2.**
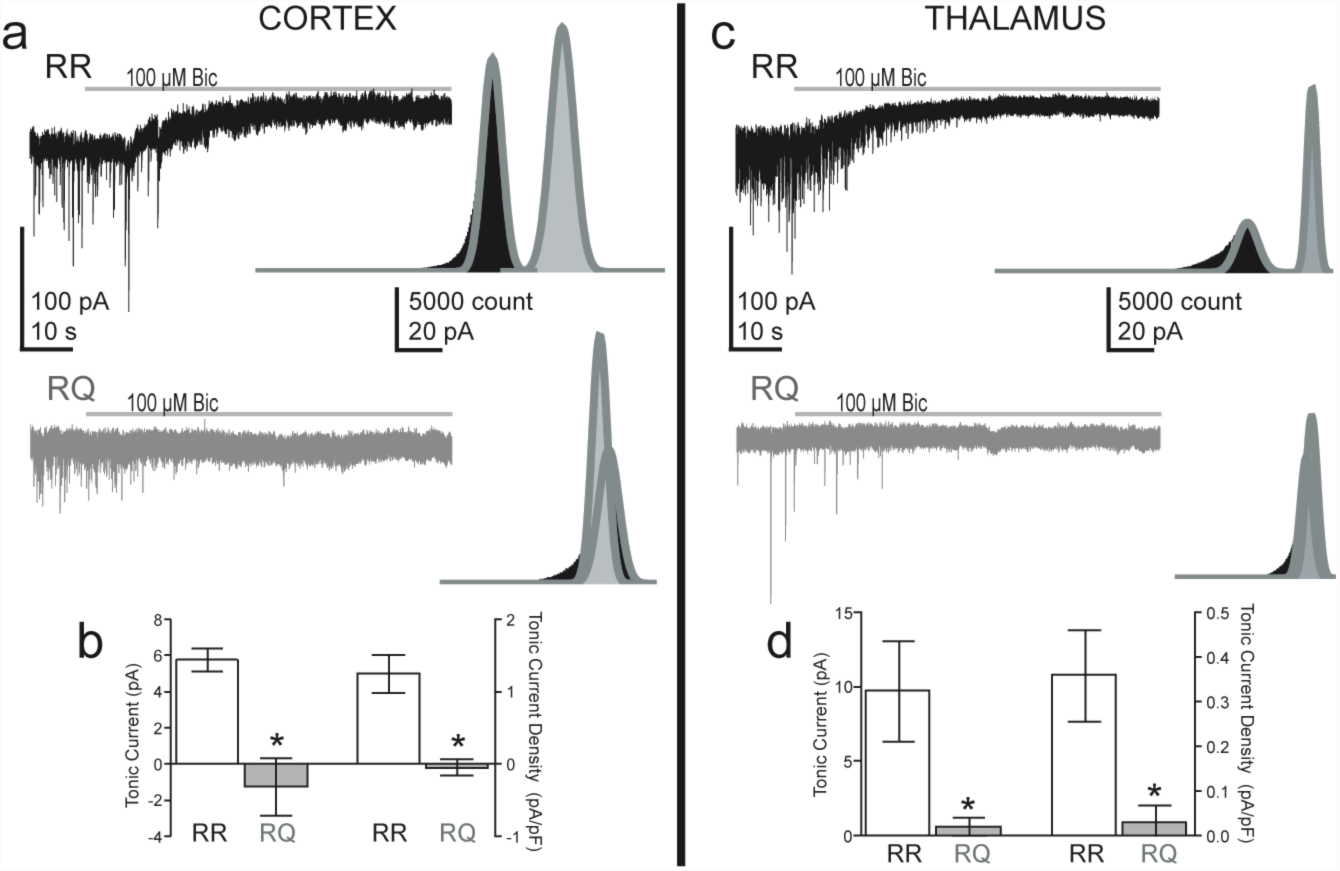
Tonic currents are abolished in RQ cortex and thalamus. A) Example voltage-clamp traces for RR (above: black) and RQ (below: grey) cortical layer II/III cell recordings during 100 μM Bicuculline administration (grey bars). Insets) Corresponding all-points amplitude histograms for data before (black) and after (grey) bicuculline administration. Histograms were fit with a Gaussian function (dark grey) only on the right side of the distribution, thus omitting components due to phasic mIPSCs. B) Tonic current amplitude (pA) (left axis) and tonic current density (pA/pF) (right axis) are abolished in RQ cortical cells (p<0.05) compared to control. C-D) Same as A-B, but for ventrobasal thalamic relay neurons.

### GABA_A_ receptor function or expression is altered in a region-specific manner

The tonic current in RR somatosensory cortical layer II/III cells (Fig. 2a) was completely blocked by the α5 subunit-selective inverse agonist L655,708 (30 μM; Fig.3a), matching previous studies showing that the α5 subunit is responsible for most or all of the native tonic inhibition in these neurons^25^. Thus the loss of cortical tonic inhibition in RQ mice may involve reduced expression or function of the α5 subunit (see below).

In contrast, application of the agonist THIP (1 μM, a concentration previously shown to be selective for δ subunit-containing receptors^20,26^) evoked currents of similar magnitude in RR and RQ cortical neurons (pA, N; RR: 21.4 ± 5.7, 4; RQ: 23.8 ± 2.2, 5; p=0.67; Fig. 3b). A similar profile of effects was observed with allopregnanolone (30 nM; Fig. 3c), a neurosteroid that also selectively activates δ subunit-containing receptors^27,28^. Together, these results suggest that receptors containing the δ subunit are present in cortical neurons, and can be recruited by both exogenous drugs and endogenous modulators, potentially providing pharmacological avenues to rescue cortical tonic inhibition in cases where it has been genetically compromised.

**Figure 3:**
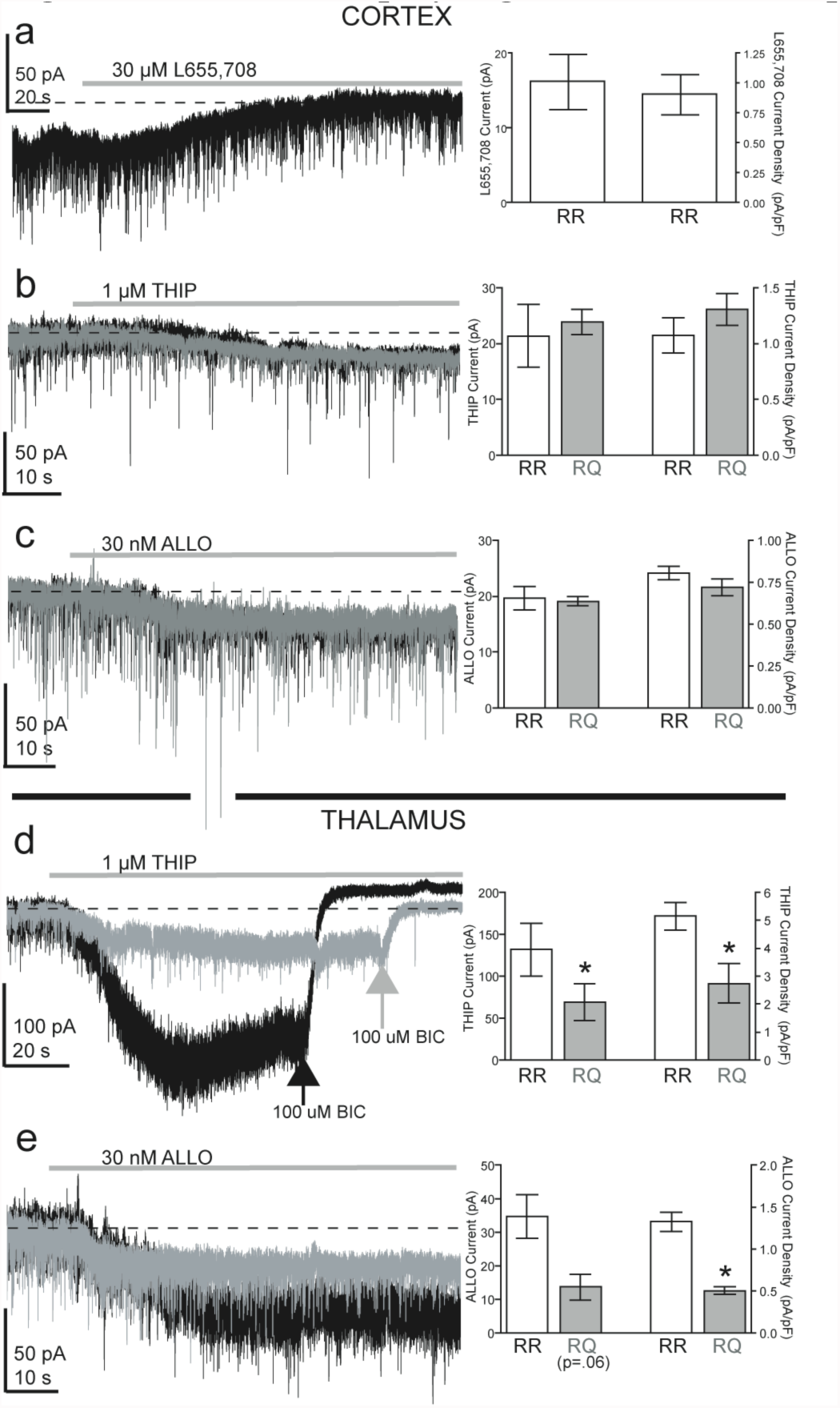
RQ mice display region- and subunit-specific changes in tonic inhibition. A) Example voltage-clamp traces for RR cortical layer II/III cell recordings during 30 μM L655,708 administration (grey bar). The current density blocked by L655,708 is not significantly different than that blocked by bicuculline (see Fig. 2). B) Both THIP (1 μM) and C) allopregnanolone (ALLO; 30 nM) induce indistinguishable current amplitude and density in RQ (grey traces) compared to RR (black traces). D) In thalamic relay neurons, however, THIP- and E) ALLO-induced current densities are significantly reduced in RQ compared to RR (~50%; p<0.05).

In contrast to cortical layer II/III neurons, thalamic relay neurons rely solely on δ subunit-containing GABA_A_ receptors to produce inhibitory tonic currents^18,20,29,30^. We found that RQ thalamic neurons responded to THIP with 47% of the current produced in RR thalamic neurons (pA, N; RR: 131.7 ± 31.2, 5; RQ: 69.3 ± 22.4, 4, p<0.05; Fig. 3d). Similarly, in RQ thalamic neurons, allopregnanolone produced 39% of the current observed in RR (pA, N; RR: 34.7 ± 6.5, 5; RQ: 13.7 ± 3.8, 3, p<0.05; Fig. 3e). These results suggest that δ subunit-containing GABA_A_ receptors are either expressed at lower levels, or have reduced activation, in thalamic relay neurons of RQ mice compared to RR mice.

### Expression of GABA_A_ receptor subunit proteins involved in tonic inhibition is reduced in RQ neurons

To test whether the loss of tonic inhibition in cortical and thalamic neurons was related to changes in the GABA_A_ receptor subunit proteins involved in tonic inhibition, we examined the levels of these proteins in whole tissue subcellular fractions (plasma membrane (PM) and intracellular organelles (ER)) using Western blotting. We calculated ‘total protein’ (PM + ER) and ‘surface trafficking’ (PM/ER) levels for all proteins assessed (α1, α4, α5, γ2, and δ). Surface trafficking was further normalized to α1-trafficking levels because previous research showed that the R43Q mutation did not alter α1 membrane trafficking^4^. We found that RQ somatosensory cortex showed a marked decrease in total α5 subunit protein expression (fraction of RR expression, N) (0.66 ± 0.14, 8, p<0.05), as well as a decrease in total γ2 subunit expression (0.76 ± 0.07, 4, p<0.05; Fig. 4b). Membrane surface trafficking was also reduced for the α4 (RQ: .59 ± .10, 8. P<0.05), α5 (RQ: 0.53 ± 0.11, 6, p<0.05), and δ (RQ: 0.60 ± 0.09, 6, p<0.05) (Fig. 4c) subunit proteins. Thus, the loss of cortical tonic inhibition we observed is consistent with the reduced expression of α5 subunits in cortical neuronal surface membranes.

**Figure 4:**
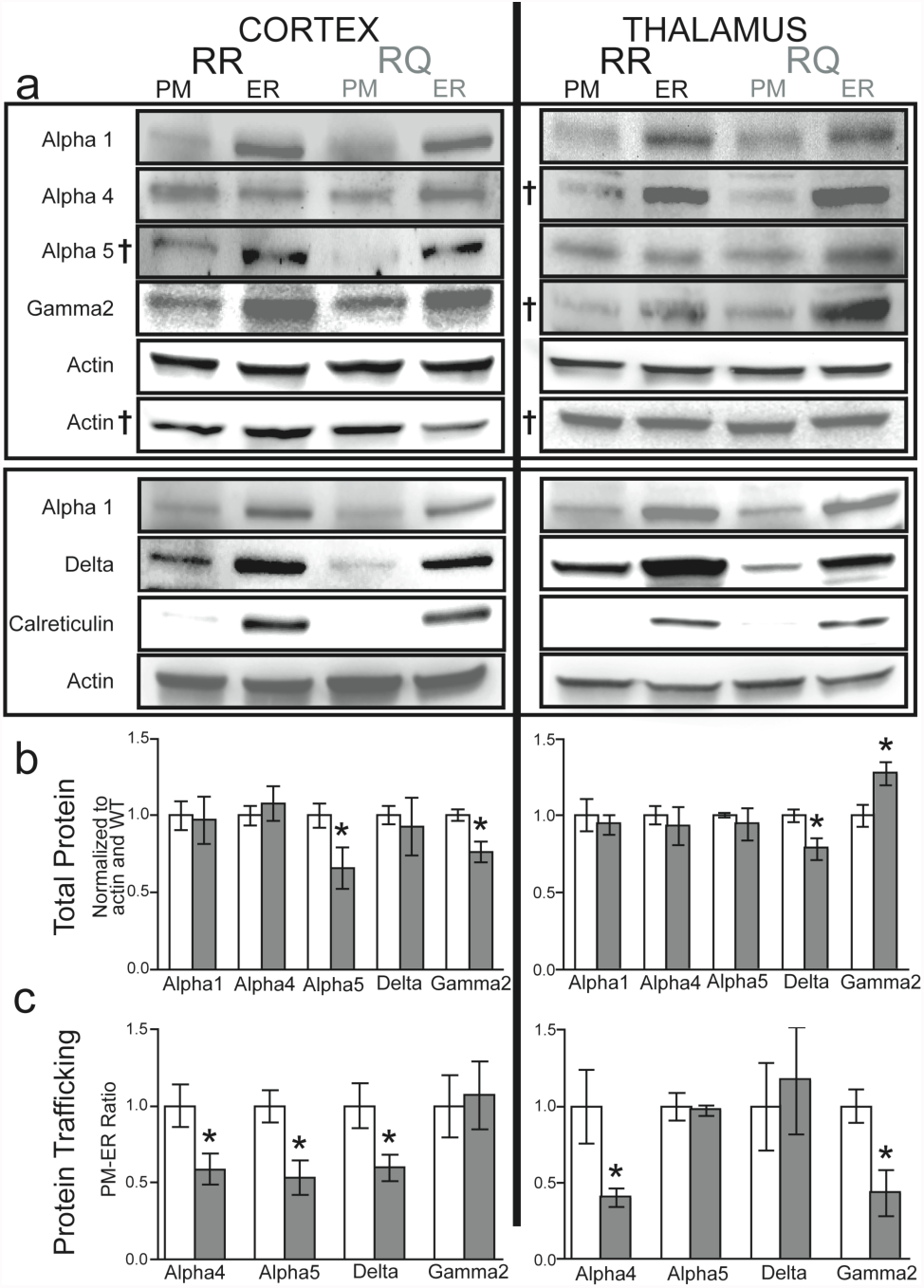
GABA_A_ receptor subunit trafficking is altered in a region-specific manner. A) Western Blots for 5 GABA_A_ receptor subunits for RR and RQ plasma membrane (PM) and intracellular organelle (ER) fractions. Calreticulin (a marker for endoplasmic reticulum) and actin were used as controls for fraction specificity and loading quantity, respectively. B) Measures of total protein (PM + ER) display decreases in α5- and γ2-subunit (p<0.05) levels in RQ cortex (grey bars) and a decrease in δ-subunit (p<0.05) in RQ thalamus compared to RR (white bars). C) Protein trafficking to the cell surface (evaluated as the PM/ER ratio) was also reduced for the α4-, α5- and δ-subunits in the cortex (p<0.05). In thalamus, trafficking to the surface was reduced for the γ2-subunit (p<0.05). Although δ subunit trafficking in the thalamus was not reduced, there was a reduction in trafficking of the α4 subunit (p<0.05), the obligatory partner for δ subunit-mediated tonic currents in thalamus.

In contrast to the cortex, thalamic tonic inhibition is mediated by receptors containing the obligatory pairing of GABA_A_ receptor α4 and δ subunits. We found that RQ thalamus showed a reduction in total δ subunit protein levels (0.78 ± 0.07, 6, p<0.05; Fig. 4B). Although we did not find a reduction in the trafficking of thalamic δ subunit protein (1.17 ± 0.35, 4, p=0.72), there was a decrease in surface trafficking of α4 subunits (0.40 ± 0.06, 5, p<0.05; Fig. 4c). Unlike RQ somatosensory cortex, RQ thalamus showed an increase in total γ2 subunit levels (1.27 ± 0.08, 6, p<0.05; Fig. 4b), but a decrease in γ2 subunit trafficking to the membrane surface (0.43 ± 0.15, 4, p<0.05; Fig. 4c).

### Firing rates and bursting behaviors are altered in RQ thalamocortical slices

Although tonic inhibition is known to contribute to neuronal responsiveness, its role in thalamocortical network activity has not been studied in detail. To explore this role, we used multielectrode extracellular recording arrays to examine neuronal spiking and burst firing in somatosensory cortex and ventrobasal thalamus of RR and RQ thalamocortical slices, as well as RR thalamocortical slices treated with L655,708 (30 μM) to selectively block α5 subunit-mediated tonic current in cortical neurons (RR-L655). Cumulative distribution plots of the average firing rates in cortical neurons show increased firing rates for RQ (median rate in Hz, [25: 75 percentiles], N; p-value (Kruskal-Wallis) (0.06, [0.02: 0.14], 230, p<0.01) and RR-L655 (0.17, [0.06: 0.22], 96, p<0.01) compared with RR (0.04, [0.01: 0.09], 328; Fig. 5c). Conversely, RQ thalamic neurons displayed decreased firing rates (0.02, [0.00: 0.13], 216, p<0.01) compared to RR (0.11, [0.03: 0.52], 444), whereas RR-L655 thalamic neurons did not (0.18, [0.02: 0.73], 122; Fig. 5c).

**Figure 5:**
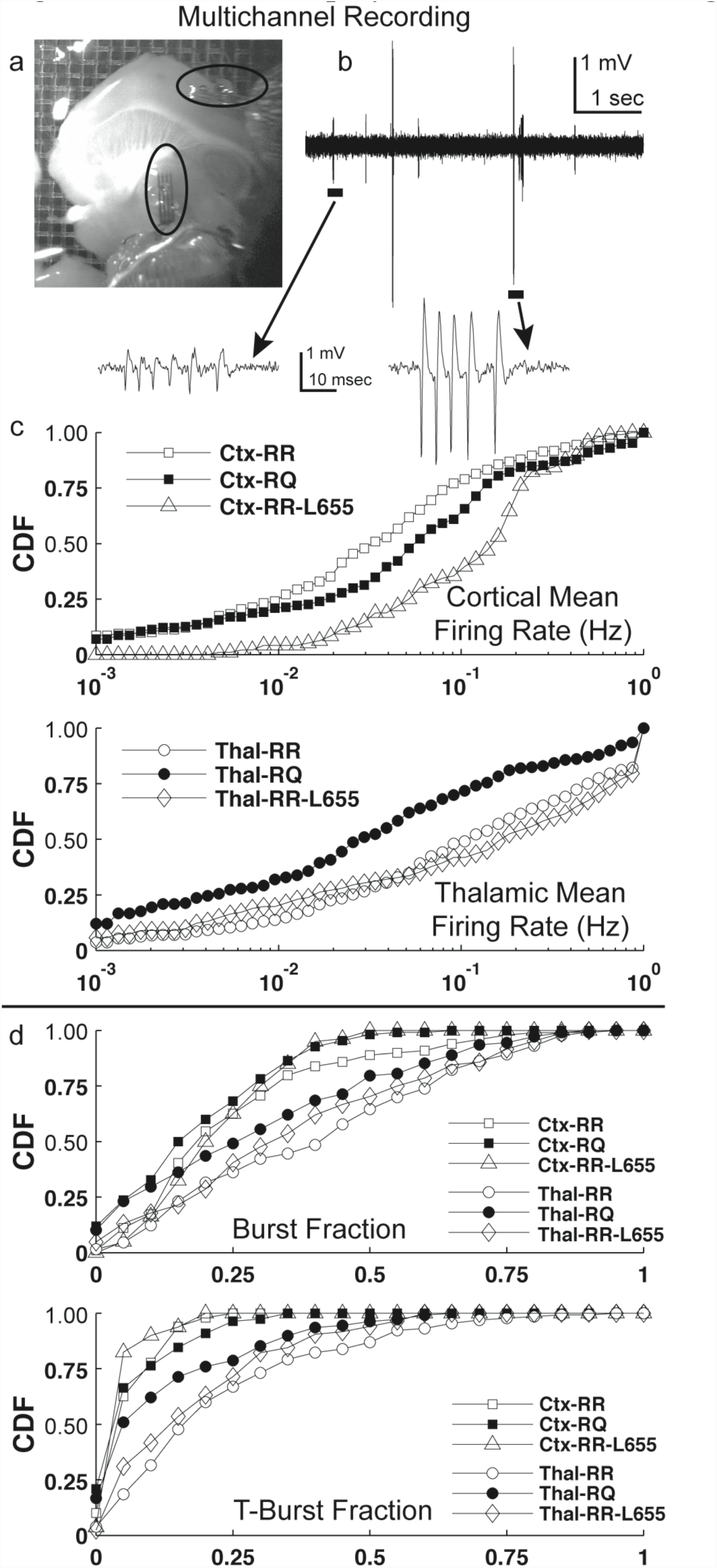
RQ slices display elevated cortical firing and reduced thalamic bursting. A) A thalamocortical slice with two multielectrode arrays (black ovals) placed in layer II/III cortex (upper) and ventrobasal thalamus (lower). B) Top: A segment of recording from an electrode located in thalamus. Bottom: Expanded segments, corresponding to the black bars in the recording above, and illustrating burst firing of two different neurons (see Methods). C) Cumulative distribution functions (CDF) of mean firing rates for cortex (CTX, upper) and thalamus (Thal, lower), for RR, RQ, and RR in the presence of L655,708 (L655). For cortex, both RQ and L655 display increased firing rates compared with RR (p<0.01). In thalamus, RQ displays reduced firing rates compared to RR (p<0.01), whereas no change is observed for RR or L655. D) CDF plots for generic burst fractions (upper) and T-burst fractions (lower, see Methods). For generic burst fraction, thalamus displayed a higher burst fraction than cortex in all conditions. In thalamus, RQ burst fraction was reduced compared to RR (p<0.05), whereas L655 was not. In cortex, neither RQ nor L655 differed from RR. For T-burst fraction, RR thalamus displayed a higher value than RR cortex (p<0.01), RQ cortex (p<0.01), and RQ thalamus (p<0.01), whereas neither RQ area was different than RR cortex.

We assessed bursting activity using two definitions of bursts: i) ‘generic’ bursts, reflecting any tendency to fire in groups of spikes, and ii) ‘T-bursts’, reflecting the temporal structure characteristic of thalamic relay neurons firing in burst mode, mediated by T-type calcium channel-dependent plateau potentials (see Methods). The ‘burst fraction’ quantified the probability that a neuron fired bursts versus lone spikes.

For generic bursts, the cortical burst fraction was lower than the thalamic burst fraction in all conditions (RR: p<0.01; RQ: p<0.05; RR-L655: p<0.05; Fig. 5d). The thalamic burst fraction was reduced in RQ compared to RR, but not in RR-L655 (RR: 0.43, [0.18: 0.64], 130; RQ: 0.29, [0.09: 0.49], 108, p<0.05; RR-L655: 0.35[0.20: 0.59], 84; Fig. 5d). There were no differences observed among cortical burst fractions (RR: 0.21, [0.14: 0.34], 99; RQ: 0.18, [0.10: 0.32], 110; RR-L655: 0.23, [0.16: 0.33], 80; Fig. 5d).

Similar to the generic burst fraction, RR cortex had a lower T-burst fraction (0.03, [0.01: 0.09], 99, p<0.01) than RR thalamus (0.16, [0.08: 0.31], 130, Fig. 5d). Addition of L655,708 to RR slices did not alter the T-burst fraction in either cortex (0.03, [0.02: 0.04], 80) or thalamus (0.15, [0.04: 0.27], 84) from control. However, the T-burst fraction was reduced in RQ thalamus compared with RR thalamus (0.05, [0.01: 0.16], 108, p < 0.01; Fig. 5d), but was not significantly different than in RQ-cortex (0.03, [0.00: 0.09], 110) compared to control.

Closer examination of T-bursts in RR slices revealed that thalamic neurons displayed more spikes per burst than cortex (cortex: 2, [2: 2], 99; thalamus: 2, [2: 3], 130; p<0.01); as well as longer burst durations (in ms; cortex: 4.1, [0.6: 5.7], 99; thalamus: 5.1, [3.6: 7.9], 130; p<0.01). Neither RQ nor RR-L655 neurons differed from control in the number of spikes per burst (RQ cortex: 2, [2: 2], 110; RQ thalamus: 2, [2: 3], 108; RR-L655 cortex: 2, [2: 2], 80; RR-L655 thalamus: 3, [2: 4], 84) or in burst durations (RQ cortex: 3.2, [0.8: 6.0], 110; RQ thalamus: 6.4, [2.6: 9.1], 108; RR-L655 cortex: 3.6, [1.5: 5.2], 80; RR-L655 thalamus: 6.1, [3.5: 8.9], 84). Similar to control, thalamus displayed more spikes per burst (p<0.01) and longer T-bursts (p<0.01) than cortex in both RQ and RR-L655.

## Discussion

Our major findings are that mice expressing the γ2R43Q mutation entirely lack GABAergic tonic currents in both somatosensory cortical layer II/III pyramidal (Fig. 2a & b) and ventrobasal thalamic relay neurons (Fig. 2c & d), and that these deficiencies increase cortical firing rates (Fig. 5c) and decrease thalamic T-bursting (Fig. 5d). The loss of tonic currents in RQ mice is correlated with decreases in surface trafficking of different GABA_A_ receptor subunits responsible for generating these currents in cortex and thalamus (Fig. 4). Selective pharmacological blockade of cortical tonic currents increased cortical firing rates as expected, but did not affect thalamic firing rates or bursting behaviors. These results are consistent with the loss of tonic currents causing neuronal depolarization that renders cortical neurons hyperexcitable and shifts thalamic relay neurons away from a burst-firing mode.

Mutation or over-expression of the γ2 subunit of the GABA_A_ receptor was previously shown to interfere with receptor assembly or trafficking^13,14,15^ of multiple GABA_A_-subunits, including the α5 subunit that mediates tonic inhibition in mouse somatosensory cortex^16,25^ and the δ subunit that mediates tonic inhibition in thalamus^18,20,29,30,31^. Our results match with these findings, showing a decrease in membrane trafficking for multiple subunits in the cortex (α5 & δ) and thalamus (α4 & γ2) (Fig. 4), and also complement previous evidence that the R43Q mutation impairs surface expression of functional GABA_A_ receptors that could result in reduced synaptic inhibition (IPSCs)^12,13,14,15,16,17^ (Fig. 1). In addition to absence epilepsy, however, this mutation also causes febrile seizures in humans and RQ mice^3,32,33^. Thus, the question arises as to whether the observed changes in tonic and phasic inhibition contribute differentially or synergistically to the absence and febrile seizure phenotypes. This issue is complicated somewhat by the variable penetrance of the absence phenotype, even amongst mice that share the C57Bl6 background^4,7,33,(present study)^, possibly due to subtle differences in genetic background between colonies or in rearing conditions. Reid and colleagues (2013) recently showed that C57Bl6 RQ mice that do not display absence seizures continue to express febrile seizures, demonstrating that the two phenotypes are dissociable in the presence of the mutation. The C57Bl6 RQ mice studied here have absence seizures (Sup. Fig. 1), have changes in both tonic and phasic inhibition and have altered thalamocortical signaling, but we have not yet measured their febrile seizure sensitivity. Thus, conclusive determination of whether the dissociation between the two phenotypes is more closely related to changes in tonic or phasic inhibition will require further research.

We showed that, in RR cortical neurons, the α5 subunit-selective inverse agonist L655,708 blocked as much tonic current as did the broad-spectrum GABA_A_ receptor antagonist bicuculline, confirming that most or all of the active tonic current in these neurons is mediated by α5 subunit-containing receptors. Thus, the loss of tonic current in RQ cortical neurons is consistent with a reduction in protein expression and trafficking of the α5 subunit in RQ, as confirmed by Western blotting (Fig. 4b & c).

We also showed that RQ thalamic neurons lack tonic currents. Furthermore, in these neurons, δ subunit-selective activators (i.e., THIP and allopregnanolone) produced less current in RQ compared to RR, suggesting dysfunction of δ subunit-containing receptors. Although Western blotting did not reveal a reduction in δ subunit surface trafficking, it did show a reduction in total δ subunit expression along with a reduction of α4 subunit trafficking, which is the partner for the δ subunit required to form functional receptors that mediate tonic inhibition in thalamic neurons^30^. Therefore, we propose that the loss of tonic inhibition in cortical and thalamic neurons in mice expressing the mutant γ2R43Q subunit is caused by a dysregulation of the assembly/trafficking of *non-mutant* subunits, namely α5 in cortex and α4 and δ in thalamus.

Reduction of inhibitory tonic currents is linked to membrane depolarization^34^, increased neuronal firing^25^ and enhanced synaptic summation^35^. Our findings that RQ somatosensory cortical layer II/III neurons lack inhibitory tonic current and exhibit increased firing rates are consistent with these previous conclusions and with the hyperexcitable cortex of human patients harboring the γ2R43Q mutation^6,7^. Although cortex and thalamus are both involved in SWDs, cortical hyperexcitability appears to be a prerequisite for SWD generation^8,9,36^, and thus the loss of cortical tonic inhibition may be a key cause of the increased intracortical excitability, increased facilitation, and the development of SWDs seen in humans harboring the γ2R43Q mutation^6^.

Thalamic relay neurons can function in either tonic or burst firing modes depending on the average membrane potential, which in turn can be influenced by the level of GABAergic tonic current^20^. Thus, depolarization resulting from the loss of tonic inhibition may shift thalamic neurons away from burst firing mode. Consistent with this idea, our multielectrode recordings revealed that RQ thalamic neurons have a reduced probability of burst firing compared with RR. Interestingly, the average thalamic firing rate was lower in RQ than in RR, suggesting that the depolarization caused by loss of tonic inhibition is relatively subtle: enough to reduce burst firing but not enough to itself promote strong tonic firing. Furthermore, selective blockade of cortical tonic inhibition with L655,708 increases the firing rate in cortex only, leaving thalamic firing and the bursting behaviors in both cortex and thalamus unaffected. Taken together, these results suggest that cortical and thalamic tonic inhibition have distinct and separable roles in regulating thalamocortical circuit function.

Previous work demonstrates a correlation between absence seizures and *enhanced* tonic inhibition in thalamic relay neurons of several rodent models^18^, leading to the conclusion that enhanced tonic GABAergic inhibition is a “necessary and sufficient condition for nonconvulsive typical absence seizure generation”^37,38^. However, our finding that γ2R43Q knock-in mice entirely lack tonic inhibition in thalamic relay neurons demonstrates that enhanced thalamic tonic inhibition is not necessary to produce absence seizures. Instead, together with the aforementioned work, our data suggest that an optimal level of tonic inhibition throughout the thalamocortical circuit is necessary for normal thalamocortical processing, such that either increases or decreases away from this optimum are sufficient to enhance susceptibility to absence epilepsy. Importantly, we also show that cortical tonic inhibition is absent and that cortical neurons have elevated firing rates in RQ mice. Future genetic or pharmacological models of region-specific deficits in tonic inhibition will be helpful for dissecting the contributions of tonic inhibition in cortex versus thalamus to regulating absence epilepsy.

Despite the absence of endogenous tonic inhibition, we show that the δ subunit-selective activators THIP and allopregnanolone can recruit tonic currents in both thalamus and cortex of RQ mice. We therefore propose that absence epilepsies can be divided into multiple classes, two distinct examples of which are characterized by either an increase^18,19^ or a decrease (e.g., γ2R43Q) in tonic inhibition. Therefore, appropriately titrated doses of tonic current activators may have high therapeutic benefit for rescuing normal function in the latter class.

## ACKNOWLEDGEMENTS

We thank Laura Ewell, Barry Schoenike, and Dan Uhlrich for their assistance and guidance of this project. This work was supported by grants from the Epilepsy Foundation (K.P.M., M.V.J.) and NIH (NS046378, NS075366 to M.V.J.).

## AUTHOR CONTRIBUTIONS

K.P.M. and M.V.J. jointly conceived and designed all experiments for the study with guidance from S.P., S.M.J., A.R., and C.C.; K.P.M., W.B.P., and A.B.N. performed experiments; K.P.M. and M.V.J. analyzed data; K.P.M and M.V.J. wrote the manuscript.

## Methods

### Whole-cell Patch Clamp Experiments

Horizontal slices (400 μm) were prepared from the brains of C57BL/6J mice of either sex (16 – 26 days old). All procedures were approved by the University of Wisconsin Institutional Animal Care and Use Committee. Mice were anesthetized with isoflurane, decapitated, and the brain was removed and placed in ice-cold cutting solution containing (in mM): 125 NaCl, 25 NaHCO_3_, 2.5 KCl, 1.25 NaH_2_PO_4_, 0.5 CaCl_2_, 3.35 MgCl_2_, 25 D-Glucose,13.87 M sucrose, and bubbled with 95% O_2_ and 5% CO_2_. Slices were cut using a vibratome (Leica VT 1000S, Global Medical Imaging; Ramsey, MN) and placed in an incubation chamber containing standard 2 mM CaCl_2_,1 mM MgCl_2_ artificial cerebrospinal fluid (ACSF) at room temperature for 1 hour before being used for recordings. Whole cell patch-clamp recordings were made from somatosensory cortical layer II/III pyramidal cells or vetrobasal thalamic relay cells, visualized using an upright differential interference contrast microscope (Axioskop FS2, Zeiss; Oberkochen, Germany). Patch pipettes were pulled from thin-walled borosilicate glass (World Precision Instruments; Sarasota, FL) with a resistance of 3-5 MΩ when filled with intracellular solution containing (in mM): 140 K-gluconate, 10 EGTA, 10 HEPES, 20 phosphocreatine, 2 Mg_2_ATP, 0.3 NaGTP (pH 7.3, 310 mOsm). Recordings were made in a submerged chamber at room temperature using a MultiClamp 700B amplifier (Axon Instruments; Foster City, CA), filtered at 4 kHz and digitized at 10 kHz using a Digidata 1322A analog-digital interface (Axon Instruments). Data were acquired to a Macintosh G4 (Apple Computer; Cupertino, CA) using Axograph X v1.1.4 (Molecular Devices; Sunnyvale, CA).

Data segments (120 s) prior to bath application of bicuculline (100 μM) were analyzed for miniature inhibitory postsynaptic currents (mIPSCs), using the variable amplitude template-matching algorithm in Axograph (τ_1_=0.64 msec, τ_2_=14.95 msec).

Additional segments (30 s) just prior to and 90 s after bicuculline administration were analyzed to quantify inhibitory tonic currents. All-point amplitude histograms were computed for each segment, and fit with a Gaussian function only to the outward current portions relative to the peak in order to omit components arising from inward phasic mIPSCs^39^. Tonic current was calculated as the difference between the fitted Gaussian means before and after bicuculline administration. Current density (pA/pF) was calculated by dividing the current by cell capacitance. Similar fitting was used to measure the currents produced by THIP (4,5,6,7-tetrahydroisoxazolo[5,4-c]pyridin-3-ol) and allopregnanolone (ALLO).

### Multichannel Electrode Array Recordings

Thalamocortical slices (400 μm)^40,41^ were prepared as above and placed on an interface chamber perfused with 3-5 ml/min of low-Mg^2+^ (200μM) ACSF. Two to four 16-channel arrays (4X4, NeuroNexus; Ann Arbor, Michigan) were inserted into somatosensory cortex or ventrobasal thalamus. Data were acquired continuously using Tucker-Davis Technologies (TDT) SH16 headstages, Medusa preamplifiers, and RX5 Pentusa Base Station (TDT; Alachua, FL) at a 12.2 kHz sampling frequency. Spikes were detected as events larger than 2.5 standard deviations above baseline noise, with 5 millisecond segments surrounding each spike captured for analysis. Spikes were sorted by principal component analysis of spike waveforms, followed by clustering of waveforms projected into the space spanned by the first three principal components using the Klustakwik algorithm^42,43^. Homewritten Matlab (MathWorks, Natick, NJ) code was used to analyze firing and bursting of each neuron, based on the timestamps of the sorted spikes. ‘Generic’ bursts were defined to reflect any tendency to fire in groups of spikes, and were detected as groups of spikes separated from other groups by gaps of ≥ 50 msec. We also used a measure specifically reflecting the expected statistics of thalamic neuron bursting mediated by T-type calcium channels (‘T-bursts’), which were detected as events with an interburst gap of ≥100 msec combined with an intraburst gap of ≤8 msec^44,45^. The ‘burst fraction’ in both cases was computed as the number of bursts containing 2 or more spikes divided by the total number of bursts.

### Subcellular Fractionation and Western Blotting

To evaluate differences in GABA_A_ receptor subunit protein expression and trafficking, somatosensory cortex and ventrobasal thalamus were dissected from horizontal slices (1200 μm) prepared as above, and immediately placed on dry ice then stored at -80° C. Samples were thawed and suspended in 50 μgL 0.1% Triton buffer with protease inhibitors (Sigma; St. Louis, MO) and further disrupted with 3-5 pumps of a fine-tipped syringe. After 10 minutes at room temperature, samples were centrifuged at 8,000*g* for 10 minutes at 4° C. The supernatant (organelle) fraction was then transferred to another chilled tube and the pellet (plasma membrane) fraction was resuspended in 50 μL Triton buffer^46,47^.

A Bradford protein assay (Bio-Rad; Hercules, CA) was performed on all samples to quantify protein concentration. Loading buffer was added, samples were boiled, and proteins were one-dimensionally separated on Mini-PROTEAN TGX (Bio-Rad) gels (10%), then transferred to polyvinylidene difluoride membranes (Immobilon®-P, Millipore; Billerica, MA). Membranes were probed with antibodies against GABA A receptor subunits α1 (#OPA1-04100: Thermo Scientific; Waltham, MA), α4 (#AB5459: Millipore), α5 (#AB9678: Millipore), δ (#868-GDN: PhosphoSolutions; Aurora, CO), and γ2 (#OPA1-04111: Thermo Scientific). Actin (#691001: MP Biomedicals; Solon, OH) and the endoplasmic reticulum-enriched protein calreticulin (#06-661: Millipore) were also probed and used as loading and organelle fraction controls, respectively^46,47^. Some gels were stripped with Restore^TM^ PLUS Western Blot Stripping Buffer (Thermo Scientific) and re-blotted for a second protein. Corresponding secondary antibodies (1:20k) (Santa Cruz Biotechnology; Santa Cruz, CA) were applied and immunolabeling of membranes was detected via SuperSignal West Femto (Thermo Scientific) chemiluminescence using a UVP ChemiDoc-IT™ Imaging System controlled by Image Acquisition and Analysis software (VisionWorks LS: UVP; Upland, CA).

### Statistics

When comparing normally distributed data, two groups were assessed with a t-test and comparisons of three or more were assessed with ANOVA. When comparing non-normally distributed data, a Kruskal-Wallis examination of medians was used to compare multiple groups.

### Supplemental Methods: RQ mice display SWDs *in vivo*

In human patients expressing the γ2R43Q mutation, penetrance of the absence epilepsy phenotype depends strongly on genetic background^3^. This dependence also applies to knock-in mice, such that penetrance can vary even between colonies of γ2R43Q mice that are nominally of the same background strain^4,7,33,49,(present study)^. Thus it is important to correlate the epilepsy phenotype with putative underlying cellular or network mechanisms. The present study used γ2R43Q knock-in mice bred into a background of Harlan C57BL/6J-OlaHsd mice. Behavioral and electrographic markers of absence epilepsy in these animals were confirmed by video-EEG monitoring. Details of surgery and electrode implantation are described in Nelson et al.^50^. Briefly, RQ mice, under isoflurane anesthesia, were implanted for chronic EEG recordings with gold plated miniature screw electrodes over the right and left frontal and parietal cortices, and one over the cerebellum as reference. Two vinyl-coated braided stainless steel wire electrodes were placed in the nuchal muscle for EMG recording. Continuous video-EEG recordings were made and SWDs were scored off-line. A SWDs event was defined as a brief (~2 seconds long) ~7 Hz signal synchronized across all EEG leads, with a corresponding lack of signal in the EMG recording. SWDs “bouts” were defined as groups of SWD events separated from other events by <1 minute.

---

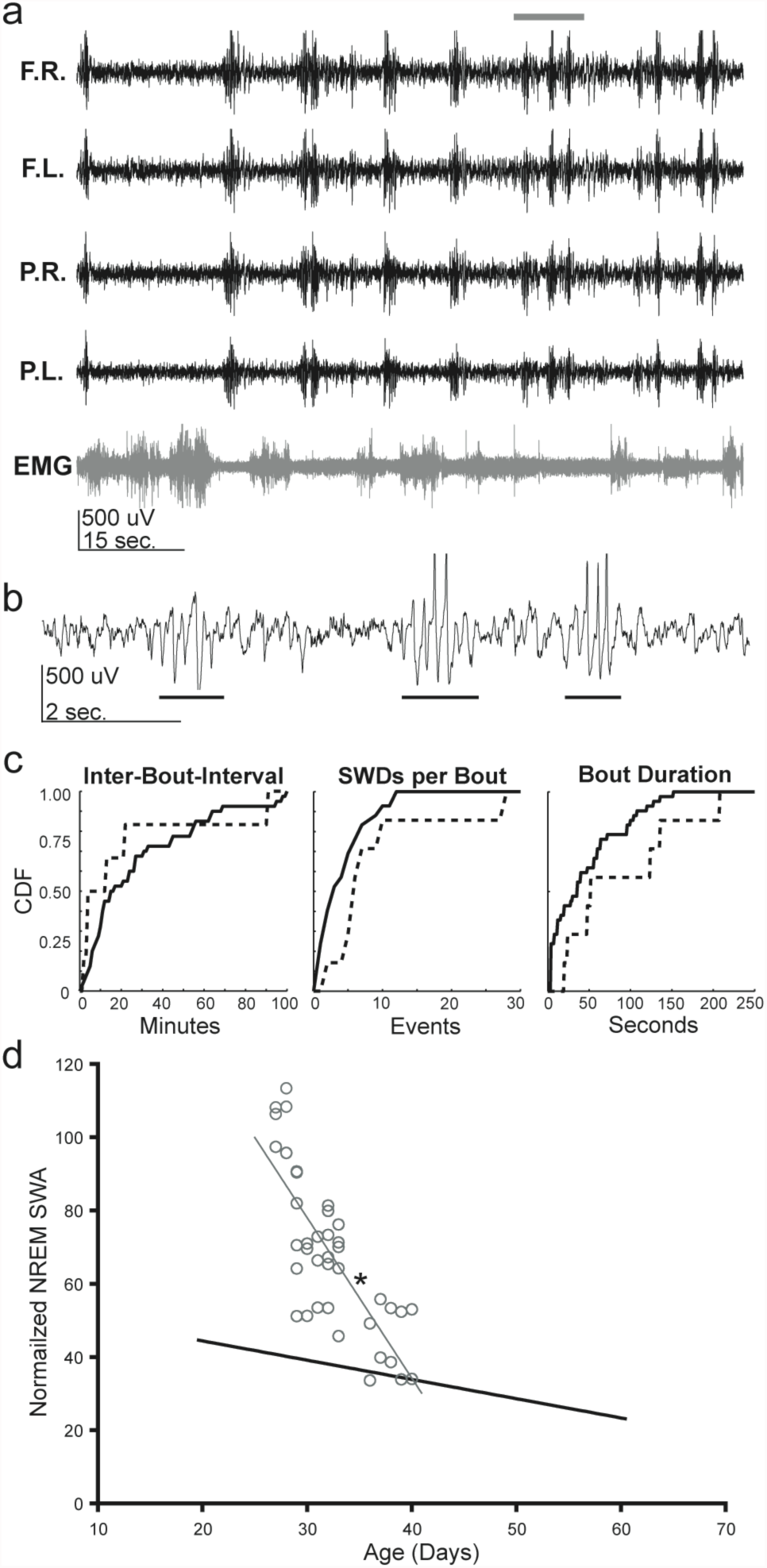
Supplemental Figure: RQ mice display SWDs *in vivo*. A) Electroencephalogram (EEG) recording of an RQ mouse. Top trace to bottom trace: frontal right cortex (F.R.); frontal left cortex (F.L.); parietal right cortex (P.R.); parietal left cortex (P.L.); electromyogram (EMG). Note the brief yet high number (~11 times during the 1.5 minute trace) of synchronized events that occur across all EEG leads during the absence of signal in the EMG. B) Expanded F.R. EEG recording from grey bar in A (10 seconds). Note the brief ~6 Hz SWD events that occur 3 times during the 10-second trace (bars). C) Cumulative distributions from two different RQ mice (solid and dashed lines) over two days of recording show similar characteristics from both animals, whereas SWDs were not observed in litter-mate control mice (not shown). D) Normalized non-rapid eye movement (NREM) slow-wave activity (SWA) power across mouse age shows that RQ mice (circles and grey line) had a steeper decline in SWA with age compared to wild-type mice (black line from ref. 46) (R^2^ = 0.682, RQ: -4.399 per day, WT: -0.526 per day; p< 0.001 by multiple regression analysis). SWA levels are an indicator of how many neurons are simultaneously entering an up-state, and thus, more cortical activity^48^.

